# THE SUBTHALAMIC NUCLEUS IS INVOLVED IN SOCIAL MEMORY IN MALE RATS

**DOI:** 10.64898/2026.06.16.732685

**Authors:** Cassandre Vielle, Lucie Vignal, Maya Williams, Nicolas Maurice, Florence Pelletier, Emilie Pecchi, Christelle Baunez

## Abstract

Human social behavior depends on complex socio-cognitive abilities, many of which are disrupted in neuropsychiatric disorders. We previously found that subthalamic nucleus (STN) lesions abolish familiarity-based modulation of social reward in rats, implicating the STN in social recognition. Here, we used pharmacological lesions, electrical deep brain stimulation (DBS), and optogenetic manipulations to dissect the STN’s role in social memory. STN lesions selectively impaired social discrimination memory without affecting social novelty discrimination or non-social memory. High-frequency optogenetic stimulation disrupted social discrimination memory only when applied during encoding, whereas STN optogenetic inhibition, and electrical DBS impaired social memory when applied during either encoding or retrieval. Notably, transient optogenetic inhibition abolished recognition of a cage-mate after short isolation, yet this ability was preserved after permanent lesions, suggesting compensatory adaptations. These results identify the STN as a key node for social memory and highlight the importance of considering its role in social cognition when implementing STN-targeted therapies.

## INTRODUCTION

The subthalamic nucleus (STN) plays a multifaceted role beyond its well-established involvement in motor control. Emerging evidence suggests its crucial contribution to cognitive and limbic functions (Baunez & Lardeux, 2011). High-frequency deep brain stimulation (HF-DBS) of the STN has emerged as a therapeutic tool for alleviating motor symptoms in Parkinson’s disease (PD) (Patricia Limousin et al., 1995; P. Limousin et al., 1995) and obsessive-compulsive disorder (OCD) (Fontaine et al., 2004; Mallet et al., 2002). Its potential extends to treating addiction (Pelloux and Baunez, 2013; Vorspan et al., 2023). However, despite these clinical applications, the intricate role of the STN in various functions remains unclear, and the implications of its neuromodulation warrant further investigation.

Several studies in rodents suggest that the STN modulates both normal and pathological social functioning (Reymann et al., 2013; Tan et al., 2011; Vielle et al., 2025; Vignal et al., 2024). Notably, in a social preference test, it has been shown that while control rats displayed greater investigative preference towards unfamiliar peers compared to cage-mates or objects, STN lesions abolished this preference, disrupting the influence of familiarity on the affective value of social interactions (Giorla et al., 2022). Additionally, while control rats self-administered less cocaine when in the presence of an unfamiliar peer than in presence of their cage-mate, STN-lesioned rats exhibited no significant difference in cocaine intake based on companion familiarity. Likewise, former work has shown that while in control rats playback of positive ultrasonic vocalizations from unfamiliar rats is more rewarding and reduces cocaine consumption compared to vocalizations from cage-mates, this difference is abolished in STN lesioned rats (Vielle et al., 2021). These findings collectively suggest that STN lesions may impair the ability of rats to differentiate familiar from unfamiliar conspecifics. This potential role in social cognition raises important considerations for patients undergoing STN HF-DBS. However, no prior studies have systematically investigated the precise role of the STN in social recognition.

Among social skills crucial for mammals living in complex social environments, like humans and rats (Barnett, 1967), social discrimination enables engagement of appropriate interactions (e.g., approach or avoidance, mating, attack) with conspecifics (Schweinfurth, 2020). The ability to discriminate and remember a peer based on past experiences is termed social memory. Social discrimination and memory are, therefore, fundamental processes in social animals. These abilities rely on specific neurobiological substrates, including the olfactory bulb (Richter, 2005), the hippocampus (Chai et al., 2021; Hitti and Siegelbaum, 2014), the medial (Ferguson et al., 2001) and basolateral amygdala (Wang et al., 2014) and some cortical areas, such as the anterior cingulate (Rudebeck et al., 2007) and the prefrontal (Tan et al., 2019) cortices. Interestingly, the STN has been shown to have structural or functional connections with these structures (Auer, 1956; Canteras et al., 1990; Çavdar et al., 2018; De Vito and Smith, 1964; Groenewegen and Berendse, 1990; Lambert et al., 2012; Monakow et al., 1978; Nambu et al., 1996; Péron et al., 2016).

Laboratory rodents can be tested for social discrimination and memory by measuring their investigatory behavior towards a novel versus a familiar peer (Engelmann et al., 1995; Thor and Holloway, 1982). When combined with optogenetic tools, such tests can help decipher the specific roles of neurobiological substrates in encoding, maintaining, and retrieving social memories (Kogan et al., 2000; Noack et al., 2015; Richter, 2005). However, a decrease in investigatory behavior could also result from deficits in novelty discrimination or general memory performance. Therefore, a battery of tests is necessary to isolate the specific process affected by the activity of a particular brain structure.

This study aimed to explore the involvement of STN in social memory. To unravel its specific role, we first assessed the effects of pharmacological lesions on rats’ performance in 1) social novelty discrimination, 2) non-social discrimination memory and 3) social memory, through a habituation-dishabituation paradigm and a social discrimination memory test. To further investigate the possible effect of STN HF-DBS on social memory in patients, we subjected other rats to electric STN HF-DBS during either the encoding or retrieval phases of the social discrimination memory test. To decipher the underlying mechanisms, we used optogenetic tools to selectively inhibit or stimulate at high-frequency STN activity during specific phases of the social discrimination memory test. Finally, we examined the effects of both short-term STN optogenetic inhibition or HF stimulation and long-term STN lesions on rats’ ability to discriminate their cage-mate from an unfamiliar peer. Through this comprehensive approach, we aim to elucidate the intricate role of the STN in social memory and its potential implications for patients undergoing STN HF-DBS.

## MATERIAL AND METHODS

### Animals

65 male Lister Hooded rats (Charles River Laboratories, Saint-Germain-sur-l’Arbresle, France) weighing ∼400 g were used in this study. Only male rats were used, because social interaction is more rewarding for them than for females (Douglas et al., 2004). Animals were housed in pairs in temperature-controlled room, maintained under a 12 h inverted light/dark cycle, with unlimited access to water and food (Scientific Animal Food and Engineering, Augy, France). Animals were handled daily for habituation. All experiments were conducted during the dark cycle (7am-7pm). All animal care and use conformed to the French regulation (Decree 2010-118) and were approved by the local ethic committee and the French Ministry of Agriculture under the License #3129.01 and followed the 3R European rules.

### Optogenetic manipulation

To transfect the STN neurons, we used AAV5 with the recombinant protein expression under CamKIIa-promoter control, obtained from UNC Vector Core (Chapel Hill, USA). The control, inhibition and HF-stimulation groups received respectively the following AAV5 constructs: AAV5-CaMKII-EYFP, AAV5-CaMKII-ArchT3.0-p2A-EYFP-WPRE and AAV5-CaMKII-hChR2(E123T/T159C)-p2A-EYFP-WPRE.

Homemade optic fibers (230µm, Thorlabs) were connected, via an optic coupler (FCMM625-50A, Thorlabs), to a 200mW 532nm DPSS laser and light pulses were generated under the control of a signal generator (DS8000 Digital Stimulator, World Precision Instruments) with 2ms light pulse at 130Hz pulse train for STN stimulation and 15s light pulse at 0.2Hz pulse train for STN inhibition. The laser power at fiber tip was set, using a power meter (PM20A, Thorlabs), at 10mW for STN HF-stimulation and 5kW for its inhibition. The effect of these light delivery parameters on STN neurons firing pattern were previously tested and validated *in vitro* and *in vivo* (Vielle et al., 2025).

### Electric HF-DBS stimulations

Homemade bilateral electrodes described in (Vignal et al., 2024) were connected, via a stimulus isolator (DLS100, WPI) and a rotating commutator (Plastic-One), to the signal generator (DS8000 Digital Stimulator, World Precision Instruments). Stimulation parameters were based on previous studies (Degoulet et al., 2021; Pelloux et al., 2018): electric pulses were generated with 80µs pulse width stimulation at 130Hz frequency. Individual stimulation intensity (50 to 150 µA) was determined for each animal just below the threshold inducing hyperkinetic movements of the contralateral paw or abnormal behaviors.

### Surgeries

All rats were anesthetized with ketamine (Imalgene, Merial, 100 mg/kg, *i.p.*) and medetomidine (Domitor, Janssen 30 mg/kg, *i.p.*), which was reversed at the end of the surgical procedure with an injection of atipamezole (Antisedan, Janssen, 0.15 mg/kg, *i.m.*). They received an antibiotic treatment with amoxicillin (Duphamox, LA, Pfizer, 100 mg/kg, *s.c.*) and were administered meloxicam (Metacam, Boehringer Ingelheim, 1 mg/kg, *s.c.*) for analgesia.

Animals were placed in the stereotaxic frame (David Kopf apparatus) and received 0.5µL bilateral injections of virus (for optogenetic rats) or 53 mM ibotenic acid (9.4 mg/mL, for STN-lesioned rats) or vehicle solution (phosphate buffer, 0.1 M; for sham-control rats) into the STN (with tooth bar set at -3.3 mm), at the following coordinates: anterior/posterior = -3.7 mm; lateral = ±2.4 mm from bregma; dorsoventral = -8.4 mm from skull; from Paxinos & Watson, 2007. For rats subjected to electric HF-DBS, electrodes were implanted into the same coordinates. For optogenetic rats, optic fibers were implanted 0.5 mm above each injection site. Electrodes and optic fibers were fixed on the skull embedded in a head-cap made of dental cement and screw on the skull.

Optogenetic et electric DBS rats were allowed 30 days for recovery from surgery and expression of the opsins. These rats were habituated to be connected to the optic coupler or the DBS device several days before the experiments. For the lesion study, sham-control and STN-lesioned rats were allowed 10 days for recovery from surgery before behavioral testing.

### Behavioral procedures

To explore the role of STN on social recognition, 6 Sham-control and 9 STN-lesioned rats were subjected to a social novelty discrimination test (1), a non-social discrimination memory test (2) and social memory testing, through a habituation-dishabituation paradigm (3) and a social discrimination memory test (4), as well as a cage-mate discrimination memory test (5). 10 electric HF-DBS rats were subjected to the social discrimination memory in three conditions: 1) with no DBS stimulation, 2) with DBS stimulation ON during the encoding phase and OFF during the retrieval phase, and 3) with DBS stimulation OFF during the encoding phase and ON during the retrieval phase.

30 optogenetic rats (10 for EYFP-control, 10 for STN inhibition and 10 for STN HF-stimulation groups) were subjected to a social discrimination memory test in two conditions 1) with the laser ON during the encoding phase and OFF during the retrieval phase, and 2) with the laser OFF during the encoding phase and ON during the retrieval phase. These optogenetic rats were also subjected to the cage-mate discrimination memory test, with the laser ON during stimuli encounter.

Animals from lesion, DBS or optogenetic groups were subjected to all conditions in a counterbalanced order. For each individual, there was a minimum interval of one day between two tests as a subject (since daily testing does not alter social investigation; Thor & Holloway, 1982). When used as a stimulus, an interval of at least four days was required before the animal could again serve as a subject.

### Apparatus

All behavioral tests were conducted in a rectangular plastic arena (40 × 30 × 40 cm; length × depth × height) containing fresh bedding, which was replaced between animals. Each animal underwent only one trial per test. Between trials, arenas and objects were cleaned with a hydrogen peroxide solution. The experimental room was kept quiet and dimly illuminated.

Each subject rat was habituated to the arena for 30 min prior to testing. Stimulus rats were age-and weight-matched to the subject rat and were habituated to handling. Different stimulus rats and objects were used for each test condition, and their assignment as “familiar” or “novel unfamiliar” stimuli was counterbalanced across subjects.

During each trial, the subject was free to explore the stimuli. Investigation time was defined, following Engelmann et al. (1995), as active exploration of a stimulus (rat or object) involving sniffing with the nose within ∼1 cm, following, nosing, grooming, or licking. All trials were recorded using a webcam connected to a computer running Bonsai software (Open Ephys). Investigatory behavior was scored offline by a trained observer.

#### 1 Social Novelty Discrimination Test

Social novelty preference was assessed using a two-trial social novelty discrimination task adapted from Engelmann et al. (1995, 2011)). During the 5-min **encoding phase**, each subject was exposed to an unfamiliar, freely moving conspecific. Immediately afterward (**recall phase**), a second novel conspecific was introduced for 5 min. Preference for social novelty was inferred when the subject spent more time investigating the novel than the familiar conspecific.

#### 2 Social Habituation–Dishabituation Test

Social memory was assessed using a habituation–dishabituation paradigm (Winslow and Camacho, 1995). Each subject rat was exposed to the same unfamiliar conspecific in four consecutive 5-min trials, separated by 10-min intertrial intervals during which the subject remained in the arena while the stimulus rat was returned to its home cage. A progressive decline in exploratory behavior across trials was interpreted as habituation. Following the fourth interval, the subject was presented with a novel unfamiliar conspecific for 5 min (**dishabituation phase**) to assess reinstatement of exploration and control for fatigue or reduced motivation.

#### 4 Social Discrimination Memory Test

Social discrimination memory was assessed using a protocol adapted from Engelmann et al. (1995, 2011). During the 5-min **encoding phase**, the subject was exposed to an unfamiliar conspecific, which was then removed for a 30-min delay. In the 5-min **recall phase**, the subject was simultaneously presented with the familiar and a novel conspecific. Intact memory was inferred from preferential investigation of the novel rat. In stimulation experiments, HF-DBS or optogenetic stimulation was delivered either during encoding, during retrieval, or not at all (stimulation-OFF). To control for handling and device connection effects, all implanted animals were connected to the stimulation apparatus regardless of stimulation status.

#### 3 Non-Social Discrimination Memory Test

Non-social object recognition memory was assessed using a protocol adapted from Engelmann et al. (1995, 2011). During the 5-min **encoding phase**, each subject was exposed to a novel object. The object was then removed for a 30-min delay. In the 5-min **recall phase**, the subject was presented with both the familiar and a novel object. Intact recognition memory was inferred from preferential exploration of the novel object.

#### 5 Cage-Mate Discrimination Memory Test

To further examine the extent of social recognition deficits, sham-operated control rats, STN-lesioned rats, and optogenetic rats underwent a cage-mate discrimination memory test. In a single 5-min **retrieval phase**, each subject was simultaneously presented with its cage-mate and a novel unfamiliar conspecific. Laser stimulation was delivered throughout the 5-min session. One STN-lesioned rat was excluded from analysis because its cage-mate died before testing.

### Immuno-histology

At the end of the experiments, sham-control, STN-lesioned and HF-DBS rats were deeply anesthetized with isoflurane and then euthanized using an intracardiac injection of pentobarbital sodium (Dolethal, Vetoquinol, 200mg/kg). Optogenetic rats were anesthetized with an overdose of a cocktail of ketamine and medetomidine (Imalgene, Merial and Domitor, Janssen 30 mg/kg, *i.p.*) and transcardially perfused with 0.1M PBS followed by 4% paraformaldehyde dissolved in PBS. Brains were removed and frozen in isopentane (Sigma-Aldrich) (after a cryoprotection in 30% sucrose for optogenetic brains) and kept at -80°C before being coronally sectioned at 40 µm thickness, using a cryostat.

Brain sections from sham-control, STN-lesioned and HF-DBS rats were stained with thionine (Sigma-Aldrich) to assess the location and extent of STN lesions (characterized by neuronal loss and associated gliosis) and the position of the electrodes implantations. Brain sections from EYFP-control and STN optogenetic inhibition rats were examined for optic fibers location and for native fluorescence expression of the opsins with an epifluorescence microscope (Zeiss, Imager.z2) immediately after being cut. In brain sections from STN HF-optogenetic stimulation animals, native fluorescence was present but required immuno-staining to control the exact boundaries of the virus expression. After PBS rehydratation (3 x 5 min), brain sections underwent a 90 min permeation step (PBS, 1% bovine serum albumin (BSA) 2% normal goat serum (NGS), 0.4% TritonX-100), PBS washes (3 x 5 min) and incubation with primary antibody (mouse anti-GFP, A11120, Life technologies; 1:200, in PBS 1% BSA, 2% NGS, 0.2% TritonX-100) at 4°C overnight. Sections were then washed with PBS (3 x 5 min), followed by 2h incubation at room temperature with secondary antibody (Goat anti-mouse Alexa 488, A11011, Life technologies, 1:400 in PBS 1% BSA, 2% NGS) and finally washed with PBS (3 x 5 min).

### Statistical Analysis

Data are presented as mean ± SEM, and statistical significance was set at α = 0.05. Analyses and graphing were performed using Prism (GraphPad Software).

During the **encoding phase**, exploration times for social stimuli were compared between groups using the Mann–Whitney U test (Sham-control vs. STN-lesioned), one-way ANOVA (EYFP-control, ArchT3.0, and hChR2 groups), or repeated-measures (RM) one-way ANOVA (HF-DBS rats).

During the **recall phase**, time spent exploring the novel versus the familiar stimulus within each group was compared using the Wilcoxon signed-rank test.

For the **social habituation–dishabituation test**, changes in exploration time across sessions were analyzed using RM one-way ANOVA, followed by Dunnett’s multiple comparisons test (session 4 as the reference), conducted separately for Sham-control and STN-lesioned rats.

## RESULTS

Representative intact and lesioned STN, as well as viral expression by EYFP-tracing fluorescence and optic fiber implantation, are illustrated in **Figure 1**. One rat was excluded from STN-lesioned group due to unsatisfying lesion (partially outside the STN). In the HF-DBS group, 2 rats were excluded due to unsatisfying placement of electrodes. In the optogenetic groups, 3 rats were excluded for absence of fluorescence in either one or both STN, and 3 others for optic fiber misplacement.

**Figure 1:**
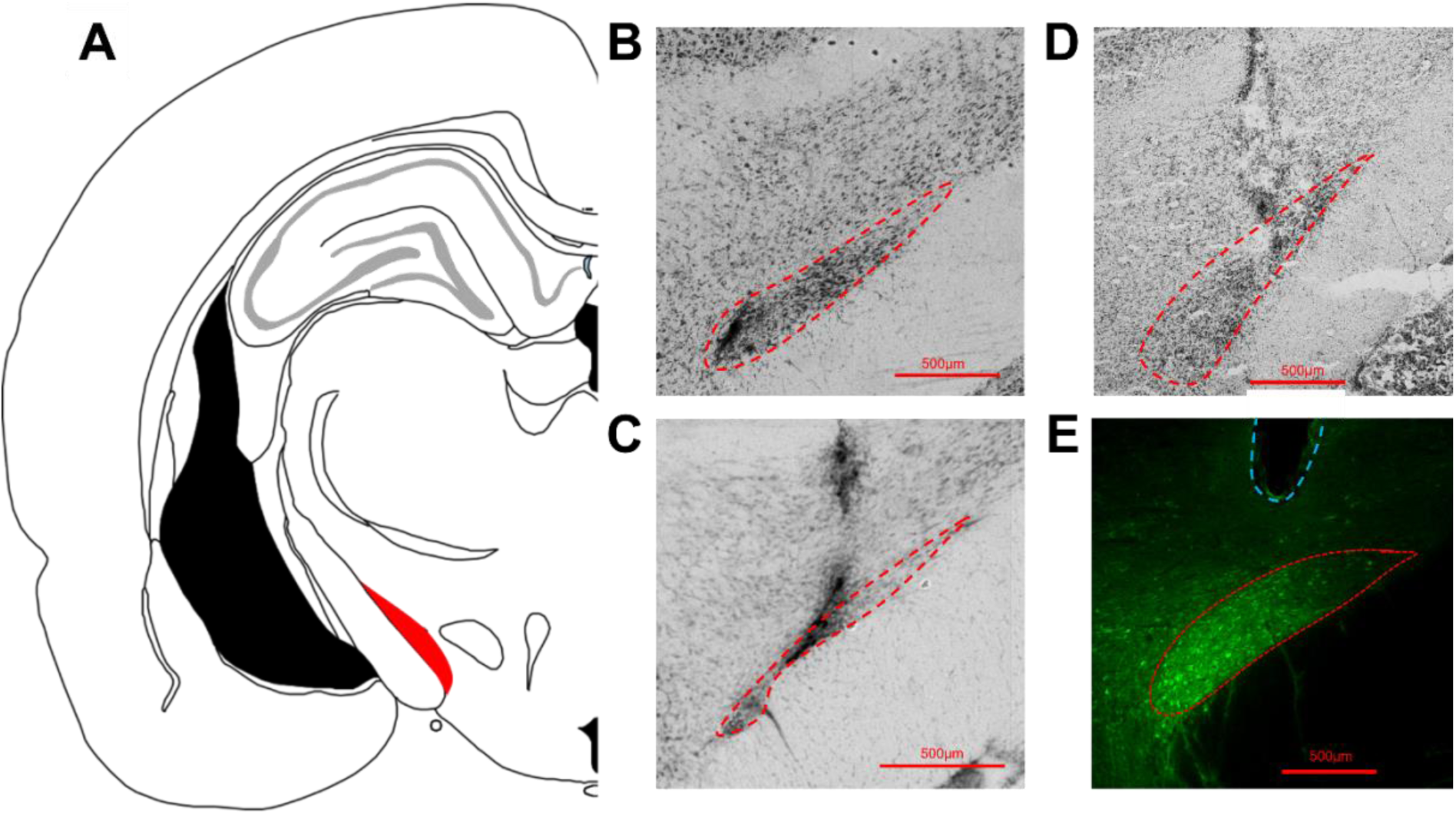
Frontal sections of the subthalamic nucleus. **A**: Schematic coronal section of the rat brain at the level of the STN (left STN in red), from Paxinos & Watson (2007). **B-D**: Representative frontal sections of the STN stained with thionine, with the STN delineated by the red lines in a **(B)** sham operated rat and **(C)** STN lesioned rat. The lesions were characterized by neuronal loss, leading to shrinkage of the structure and gliosis. **(D)** DBS-operated rat, showing a mechanical lesion caused by electrode wires. **(E)** Representative native fluorescence emitted by the transfected STN neurons (delineated by the red dotted line) and implanted optic fiber trace (delimited by the blue dotted line).

### STN lesions do not affect social novelty discrimination

To assess the effect of STN lesions on social novelty discrimination, sham-control (n=6) and STN-lesioned (n=8) rats underwent a social novelty discrimination test (**Fig 2A**). During the encoding phase, both groups spent a comparable amount of time exploring the social stimulus (**Fig. 2B**; sham-control: 103±17 s, STN-lesioned: 118±13 s; U=19, p=0.573). In the recall phase, both sham and STN-lesioned rats spent significantly more time exploring the novel compared to the familiar stimulus (**Fig. 2C-D**; sham-control group: W=21, p<0.05; STN-lesioned group: W=36, p<0.01). The mean time spent, respectively by sham-control and STN lesioned rats, exploring the novel: 70±7s, 68±6 s and the familiar peer: 34±4s, 39±5s indicates that STN lesions do not affect social novelty discrimination.

**Figure 2:**
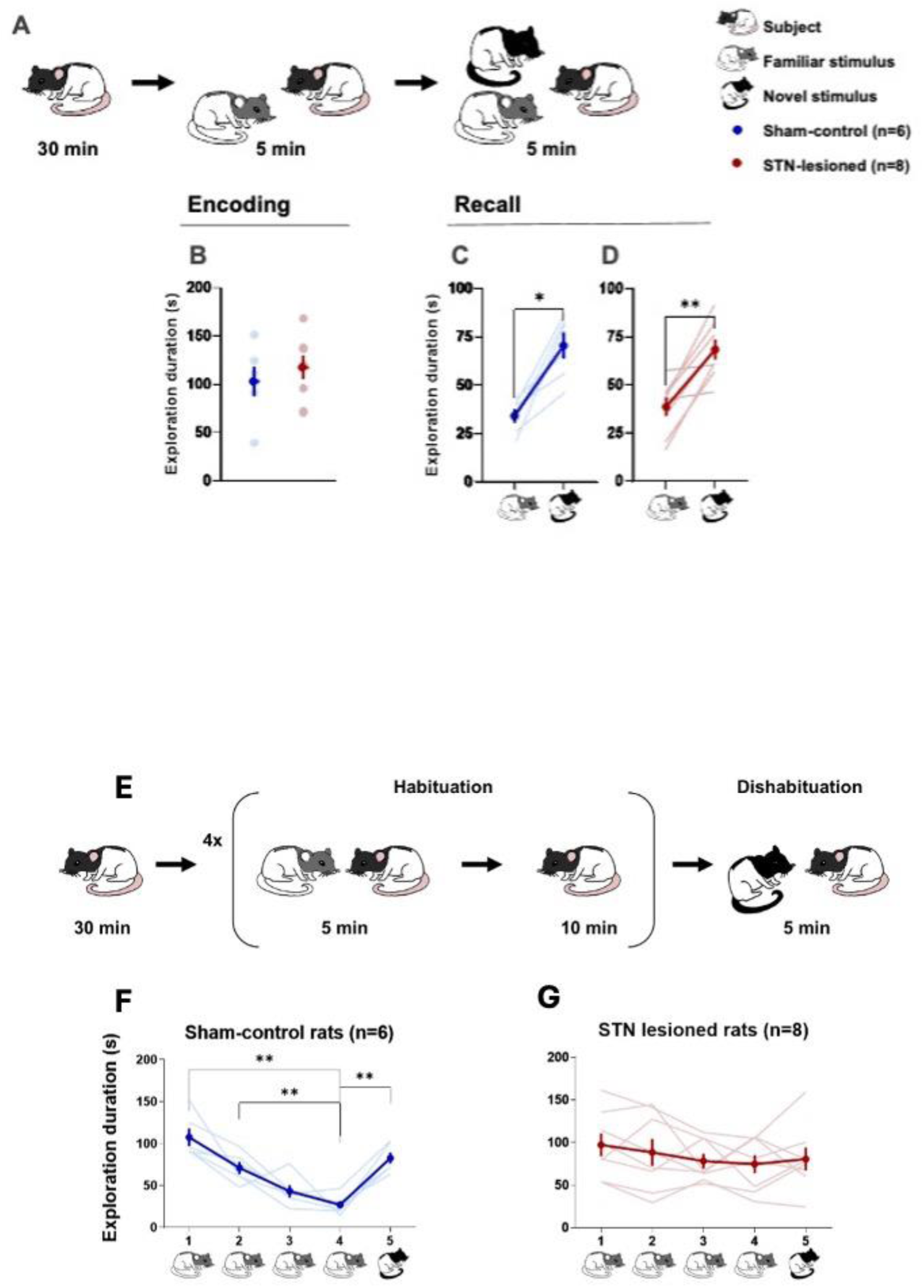
STN lesions do not affect social novelty discrimination, but impair social habituation-dishabituation. **A**. Schematic representation of the procedure to test social novelty. After a 30-min habituation to the arena (Left), the subject rat was exposed to a social stimulus (becoming the familiar stimulus at recall, grey and white) for 5-min (encoding phase). <u>Immediately after</u>, a novel social stimulus (black and white) was introduced in the arena for an extra 5min (recall phase). **B.** Mean time (±SEM) spent by the sham-control (n=6, in blue) and STN-lesioned (n=8, in red) rats exploring the social stimulus during the encoding phase. **C, D.** Mean time (±SEM) spent by the sham-control (C) and STN-lesioned (D) rats exploring the familiar (left) and novel (right) stimulus rat during the recall phase. *: p<0.05 and ***: p<0.01 *familiar vs. novel stimulus.* The dim circles and lines illustrate individual data. **E.** Schematic representation of the procedure to test social habituation-dishabituation. After a 30-min habituation to the arena, the subject rat is submitted to the habituation phase that consists in 4 encounters of 5-min with the same stimulus animal (in grey and white), separated by 10-min intervals. Then, it is exposed to the dishabituation phase that consists in 5-min encounter with a novel rat (in black and white). **F,G.** Mean time (±SEM) spent by the sham-control (F; blue dots and lines) and STN-lesioned (G; red dots and lines) rats exploring the familiar (grey) social stimulus during the 4 encounters of the habituation phase and the novel (black and white) one during the dishabituation phase (encounter 5). **: p<0.01, *vs. encounter 4.* The dim lines represent individual data.

### STN lesions impair social habituation-dishabituation

We next assessed short-term social recognition using a habituation-dishabituation protocol in sham-control (n=6) and STN-lesioned (n=8) rats (**Fig 2E**). Sham rats showed a progressive decrease in exploration over repeated exposures to the same stimulus (habituation phase; encounter 1-4; (**Fig. 2F**; repeated-measures ANOVA: F_(2.698,13.49)_ = 15.61, p = 0.0001; multiple comparisons test: encounter 4 vs.1 mean difference -80.45 (95%CI: -123.2 to -37.67), p=0.0037, encounter 4 vs. 2 mean difference: -43.91(95%CI: -71.43 to -16.39), p=0.008), followed by a marked increase when presented with a novel conspecific in the dishabituation phase (encounter 5; encounter 5 vs. 4: mean difference= -55.68 (95%CI: -80.62 to -30.74), p=0.002). In contrast, STN-lesioned rats did not show any significant change in exploration across trials (**Fig 2G**; F_(2.858,20)_=1.414, p=0.2682), revealing that STN lesions impair social habituation-dishabituation.

### STN lesions disrupt social discrimination memory

To evaluate longer-term social memory, sham-control (n=6) and STN-lesioned (n=8) rats were tested in a social discrimination memory test (**Fig 3A**). Both groups explored the social stimulus similarly during the encoding phase (**Fig 3B**; sham-control: 97±9s and STN-lesioned animals: 96±10s; U=20, p=0.662). In the recall phase, sham rats preferred the novel peer (78±13s) than the familiar one (45±8s) (**Fig 3C**; W=21, p=0.031), whereas STN-lesioned rats did not show any preference (novel: 50±8s, familiar: 51±7 s; **Fig. 3D**; W=0, p>0.999), indicating a deficit in social discrimination memory.

**Figure 3:**
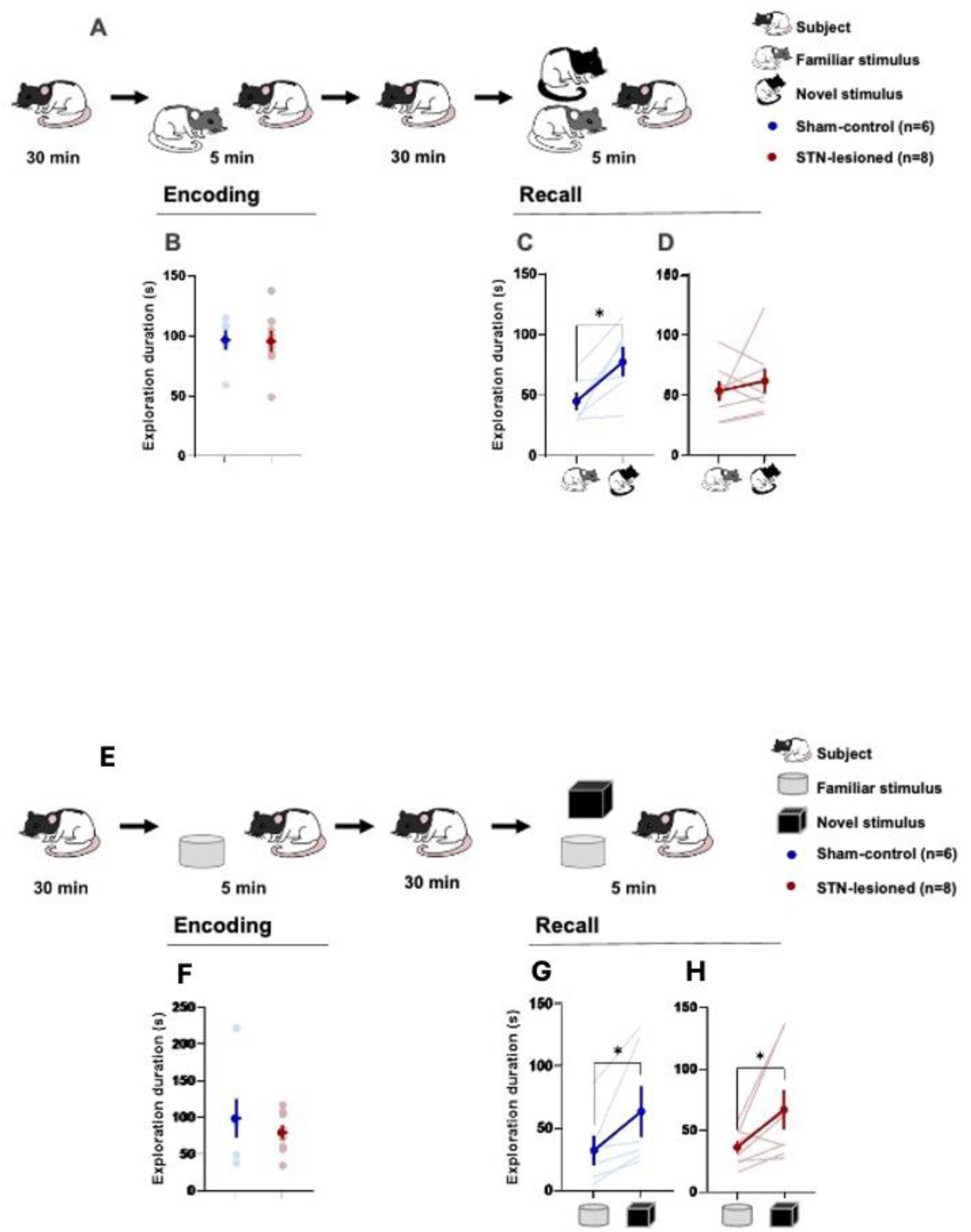
STN lesions disrupt social discrimination memory, but do not affect non-social one. **A.** Schematic representation of the procedure for social discrimination memory. After a 30-min habituation to the arena (Left), the subject rat was exposed to a social stimulus (becoming the familiar stimulus at recall, grey and white) for 5-min (encoding phase). <u>After a 30-min intertrial</u> <u>interval</u>, the subject animal was re-exposed to the same peer (familiar stimulus, in grey) and a novel rat (black and white), for an extra 5min (recall phase). **B.** Mean time (±SEM) spent by the sham-control (n=6, in blue) and STN-lesioned (n=8, in red) rats exploring the social stimulus during the encoding phase. **C, D.** Mean time (±SEM) spent by the sham-control (C) and STN-lesioned (D) rats exploring the familiar (left) and novel (right) peers during the recall phase. B-D: The bright circles and lines illustrate individual data. **E.** Schematic representation of the procedure for non-social discrimination memory. After a 30-min habituation to the arena (Left), the subject rat was exposed to an object (becoming the familiar stimulus at recall, grey) for 5-min (encoding phase). After a 30-min intertrial interval, the subject animal was re-exposed to the same object (familiar stimulus, in grey) and a novel object (black), for an extra 5min (recall phase). **F.** Mean time (±SEM) spent by the sham-control (n=6, in blue) and STN-lesioned (n=8, in red) rats exploring the object stimulus during the encoding phase. **G, H.** Mean time (±SEM) spent by the sham-control (C) and STN-lesioned (D) rats exploring the familiar (left; grey round) and novel (right; black cube) objects during the recall phase. F-H: The dim circles and lines illustrate individual data. *: p<0.05 *familiar vs. novel stimulus*.

### STN lesions do not affect non-social discrimination memory

To control for non-social memory, sham-control (n=6) and STN-lesioned (n=8) rats were then subjected to a non-social discrimination test (**Fig 3E**). Both groups showed similar exploration during the exploring/encoding phase (**Fig 3F**, sham-control: 99±29s, STN-lesioned animals: 80±11s; U=23, p=0.9497). During the recall phase, sham-control rats spent more time exploring the novel object (63±22s) than the familiar one (32±13s) (**Fig 3G**; W=21, p=0.031). STN-lesioned rats also preferred the novel object (67±17s) over the familiar one (36±6s) (**Fig 3H**; W=32, p=0.023), showing that non-social discrimination memory is preserved in STN-lesioned rats.

### STN HF-DBS alters social discrimination memory when applied at either encoding or retrieval

Since lesions are permanent at each phase, we used STN high-frequency deep brain stimulation (HF-DBS) during either encoding or recall in the social discrimination memory test in rats (n=8) (**Fig 4A**). Social exploration during encoding was unaffected by stimulation (**Fig 4B**; RM one-way ANOVA stimulation condition: F_(1.318,9.223)_=0.5851, p=0.509; exploration duration without stimulation (0-0 Hz) 52±14s, stimulation applied only during the encoding phase (130-0 Hz) 66±17s, stimulation applied only during the recall phase (0-130 Hz) 60±11s). In absence of any stimulation (0-0Hz), rats showed a clear preference for the novel peer (37±5s) over the familiar one (21±3s; **Fig. 4C**; W = 36, p = 0.008) during the recall phase. However, when stimulation was applied only during encoding (**Fig. 4D**), exploration was equivalent for both stimuli at recall (novel: 32±4s, familiar: 27±6s; W = 4, p = 0.844). Similarly, stimulation applied only during recall abolished discrimination (**Fig 4E**; novel: 32±2s, familiar: 30±5s; W=16, p=0.3125), indicating that HF-DBS applied either at encoding or retrieval affect recall and thus social memory.

**Figure 4:**
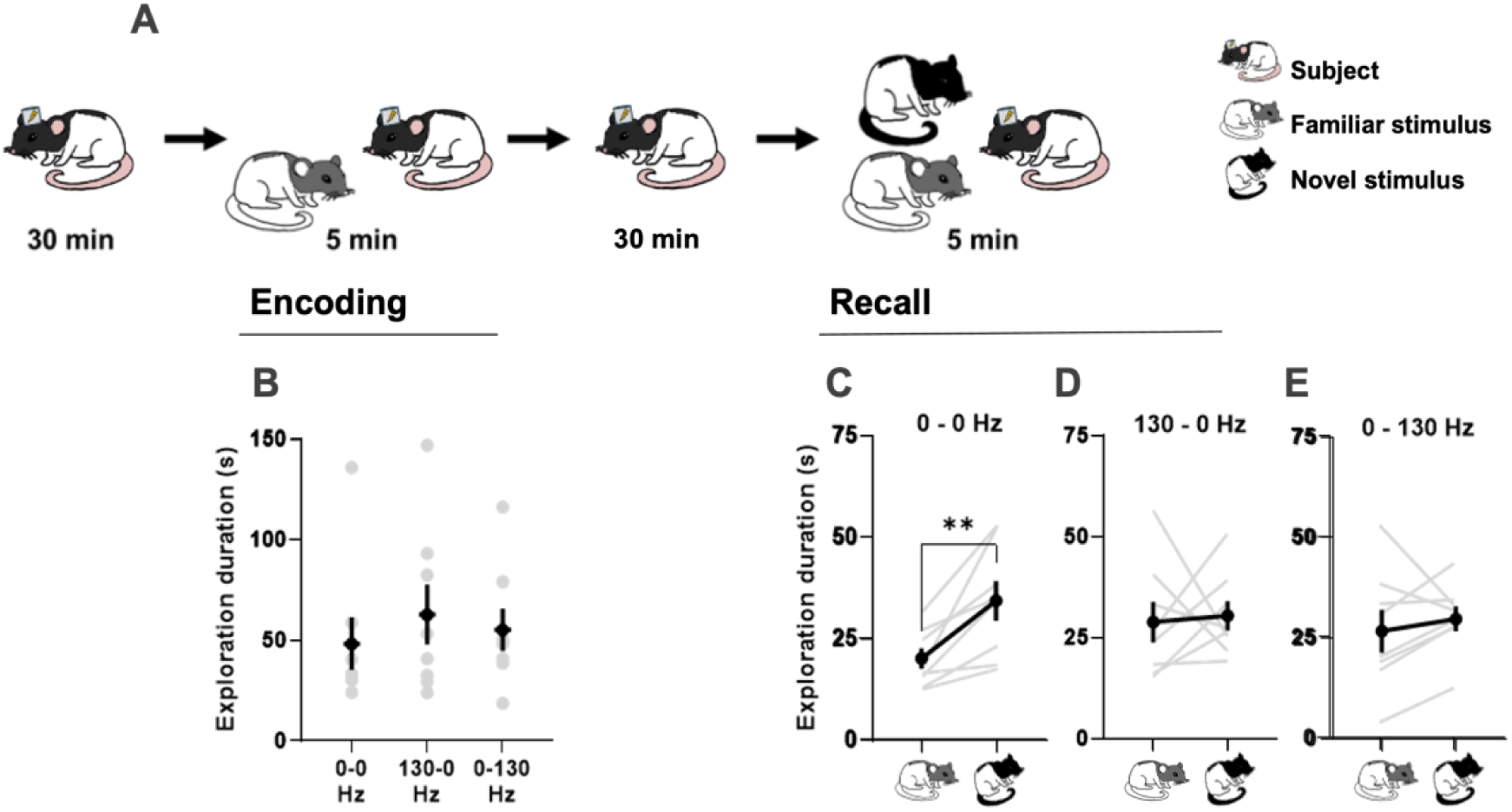
STN HF-DBS alters social memory discrimination. **A.** Schematic representation of the procedure. After a 30-min habituation to the arena (Left), the subject rat was exposed to a social stimulus (familiar stimulus, grey and white) for 5-min (encoding phase). After a 30-min intertrial interval, the subject animal was re-exposed to the same peer (familiar stimulus, in grey) and a novel rat (black and white), for an extra 5min (recall phase). **B.** Mean time (±SEM) spent by the animals (n=8) exploring the social stimulus during the encoding phase, when no stimulation was applied (0-0 Hz), when the stimulation was applied during the encoding only (130-0 Hz) or the recall only (0-130 Hz) phase. **C, D, E.** Mean time (±SEM) spent by rats exploring the familiar (left) and novel (right) peers during the recall phase, when no stimulation was applied (C; (0-0 Hz)), when the stimulation was applied only during the encoding(D; 130-0 Hz)), or the decoding (E; 0-130 Hz) phase. **: p<0.01 *familiar vs. novel stimulus.* The dim circles and lines illustrate individual data.

### STN photo-inhibition applied at either encoding or retrieval impairs social discrimination memory, while only photo-stimulation applied at encoding does

Since it is still not clear whether STN DBS induces an inhibition of STN neurons, we used optogenetic inhibition (ARCHT3.0), high-frequency photo-stimulation (hChR2) and EYFP-control manipulations applied either during encoding or recall of the social discrimination memory test (**Fig 5A**). During encoding with laser ON, exploration was similar across groups (**Fig 5B**; one-way ANOVA F_(2,21)_=1.804, p=0.1893; EYFP-control 74±8s, ARCHT3.0 93±11s and hChR2 rats 69±11s). In the recall phase (laser OFF), EYFP-controls spent more time with the novel peer (48±5s) than the familiar one (24±4s) (**Fig 5C**, W=55, p=0.002), while ARCHT3.0 and hChR2 rats showed no preference (ARCHT3.0: novel 48±5s, familiar 39±5s; **Fig. 5D**; W=7, p=0.564; hChR2: novel 44±6s, familiar 40±4s; **Fig. 5E**; W=5, p=0.688), indicating that intact STN is necessary at encoding. When the laser was ON only during recall, the exploration during the encoding phase remained unaffected with no differences between groups (**Fig 5F**; one-way ANOVA F_(2,21)_=1.416, p=0.265; EYFP-control: 83±10s, ARCHT3.0: 91±7s and hChR2 rats: 68±11s). At recall, EYFP-controls and hChR2 rats preferred the novel peer (EYFP: novel 57±5s, familiar 37±8s; **Fig. 5G**; W=39, p=0.049; hChR2: novel 34±5s, familiar 20±3s; **Fig. 5I**; W=24, p=0.0469), whereas ARCHT3.0 rats did not (novel: 42±6s, familiar: 47±9s; **Fig 5H**, W=-7, p=0.813), indicating that only STN inhibition, but not high frequency stimulation affects retrieval at this stage.

**Figure 5:**
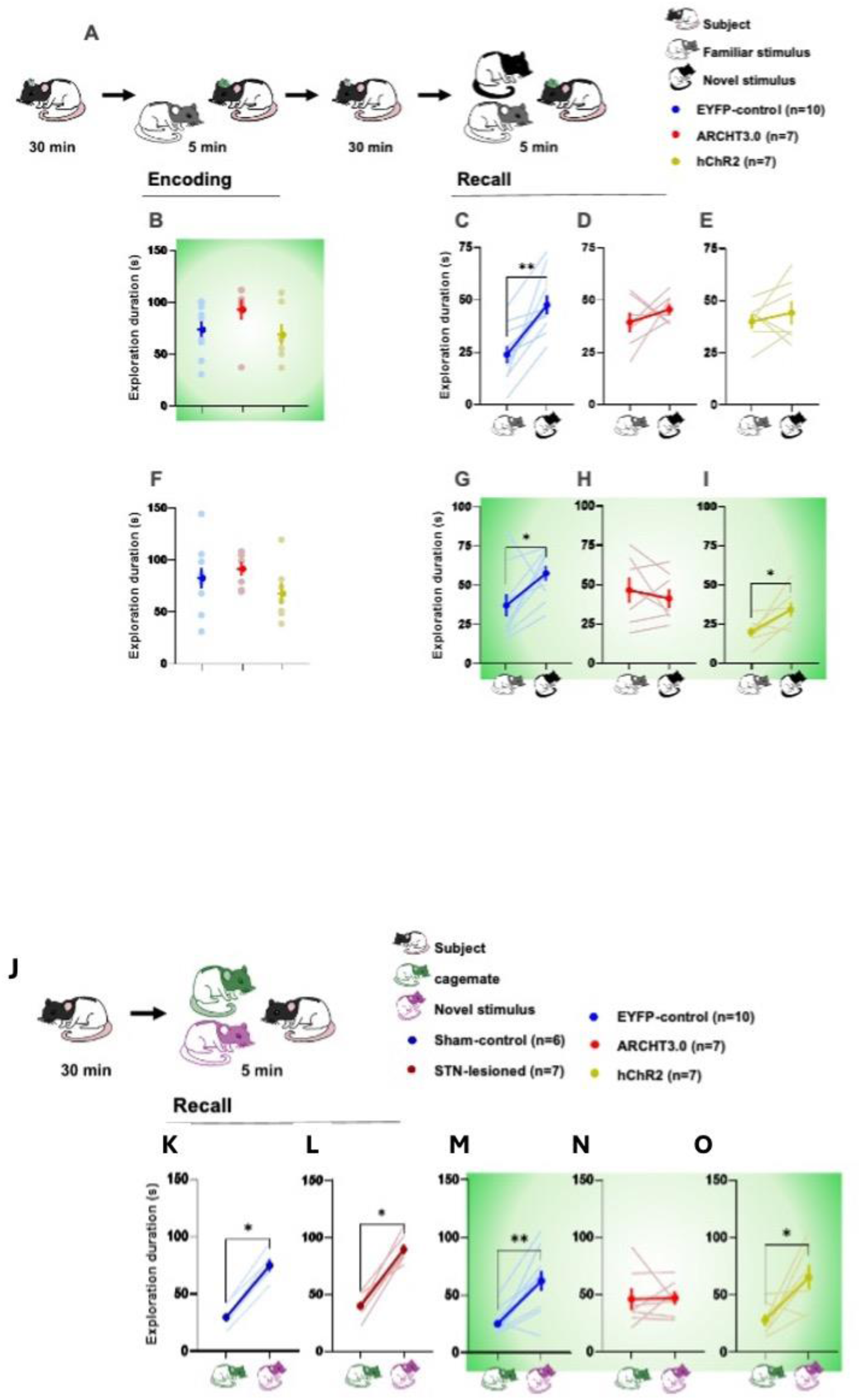
A-I: STN photo-inhibition applied at either encoding or retrieval impairs social discrimination memory, while only photo-stimulation applied at encoding does. 5J-O: STN photo-inhibition impairs cage-mate discrimination memory, unlike lesions or HF photo-stimulation. **A.** Schematic representation of the procedure to test social discrimination memory with optogenetic manipulations. After a 30-min habituation to the arena (Left), the subject rat was exposed to a social stimulus (familiar stimulus, grey and white) for 5-min (encoding phase). After a 30-min intertrial interval, the subject animal was re-exposed to the same peer (familiar stimulus, in grey) and a novel rat (black and white), for an extra 5min (recall phase). The laser activation (in green) took place during the encoding (B-E) or the recall (F-I) phase of the test. **B.** Mean time (±SEM) spent by the EYFP-control (n=10, blue), ARCHT3.0 (n=7, red) and hChR2 (n=7, yellow) exploring the social stimulus during the encoding phase, when the laser stimulation was applied. **C, D, E.** Mean time (±SEM) spent by rats exploring the familiar (left) and novel (right) peers during the recall phase, when no stimulation was applied for the EYFP-control (C), ARCHT3.0 (D) and hChR2 (E) rats. **F.** Mean time (±SEM) spent by the EYFP-control (n=10, blue), ARCHT3.0 (n=7, red) and hChR2 (n=7, yellow) exploring the social stimulus during the encoding phase, when no laser stimulation was applied. **G, H, I.** Mean time (±SEM) spent by rats exploring the familiar (left) and novel (right) peers during the recall phase, when the laser stimulation was applied for the EYFP-control (G), ARCHT3.0 (H) and hChR2 (I) rats. *: p=0.05, **: p<0.01 *familiar vs. novel stimulus.* The dim circles and lines illustrate individual data. **E.** Schematic representation of the procedure to test cage-mate discrimination memory. After a 30-min habituation to the arena (Left), the subject rat was exposed to its cage-mate (in green) and a novel rat (in purple), for 5min (recall phase). **K, L, M, N, O.** Mean time (±SEM) spent by rats exploring the cage-mate (left) and novel peer (right) during the recall phase by Sham-control (K) and STN-lesioned rats (L), and by optogenetic rats subjected to the laser stimulation (in green; M-O): EYFP-control (M), ARCHT3.0 (N) and hChR2 (O). *: p<0.05 and **: p<0.01 *cage-mate vs. novel stimulus.* The bright circles and lines illustrate individual data.

### STN photo-inhibition impairs cage-mate discrimination memory, unlike lesions or HF photo-stimulation

Finally, to manipulate the familiarity to a deeper level, we used a cage-mate discrimination memory test in which sham-control, STN-lesioned and optogenetic rats were tested (**Fig 5J**). Sham-control rats (n=6) explored the novel peer more (75±6) than their cage-mate (30±4s) (**Fig 5K**; W=21, p=0.031). STN-lesioned rats (n7) showed similar discrimination (novel: 90±5s, cage-mate: 40±4s; **Fig 5L**; W=28, p=0.016), as did EYFP-control rats (n=10) (novel: 63±9, cage-mate: 26±4s; **Fig 5M**; W=53, p=0.004). In contrast, ARCHT3.0 rats (n=7) failed to discriminate the novel rat from their cage-mate (novel: 47±6s, cage-mate: 46±10; **Fig 5N**; W=2, p=0.938), while hChR2 rats (n=7) retained discrimination (novel: 65±12, cage-mate: 28±6s; **Fig 5O**; W=24, p=0.047). These findings indicate that selective STN inhibition impairs retrieval of cage-mate memory, while lesions and high frequency photo-stimulation do not.

## DISCUSSION

Our results demonstrate that the STN plays a critical role in social memory processes that relies on familiarity and duration. Indeed STN lesioned or manipulated rats showed similar levels of social exploration during initial encounters and lesioned rats could discriminate a novel peer immediately encountered after interaction of another peer, but failed if a 30 minute interval was introduced.

STN-lesioned rats failed also to recognize a previously encountered peer in the habituation-dishabituation paradigms. This impairment cannot be attributed to general memory, as STN-lesioned rats performed normally in object discrimination memory. Interestingly, STN-lesioned rats were able to recognize their cage-mate after 30-min of separation, but not specific optogenetic inhibition of STN neurons. Alteration of the STN functioning at either encoding or recall affected the ability to discriminate between a novel and a previously encountered peer, suggesting that the integrity of STN functioning is necessary to be able to memorize social information regarding the identity of a peer.

Our study provides the first direct evidence implicating the STN in social recognition memory. Previous research has shown that social stimuli, such as the presence of a peer or playback of ultrasonic vocalizations, are modulated by familiarity and carry rewarding value (Giorla et al., 2022; Vielle et al., 2021). In those studies, STN lesions abolished the differential reward value of familiar versus unfamiliar peers or ultrasonic vocalizations. The discrepancy with our results, where cage-mate recognition was preserved, may stem from methodological differences. In our experiments, rats had unrestricted access to conspecifics, including tactile and non-volatile olfactory cues (Engelmann et al., 2011), which are critical for individual recognition in rodents. In contrast, in Giorla et al. (2022) and Vielle et al. (2021), social stimuli were restricted behind barriers or reduced to acoustic signals, limiting access to such cues and likely exacerbating recognition deficits in STN-lesioned rats.

These findings have potential implications for understanding social behavior disturbances in patients undergoing STN deep brain stimulation (DBS), particularly for Parkinson’s disease (PD) or obsessive-compulsive disorder (OCD). While impairments in emotion recognition are well documented in these patients (Biseul et al., 2005; Brück et al., 2011; Kalampokini et al., 2020; Péron et al., 2010b), deficits in face recognition are typically absent (Biseul et al., 2005; Drapier et al., 2008; Péron et al., 2010a). This apparent discrepancy could reflect differences in sensory modalities and memory systems being recruited. Rodent social recognition relies heavily on olfaction and may reflect “covert” memory processes—unconscious and affectively driven—whereas human studies generally assess explicit, “overt” memory, often restricted to visual information (Tranel and Damasio, 1993, 1990, 1988, 1987), despite the recently highlighted contribution of the olfactory system in human cognition (Endevelt-Shapira et al., 2018; Frumin et al., 2015; McGann, 2017). Given the STN’s known role in emotional salience (Pelloux et al., 2014; Péron et al., 2013; Serranová et al., 2011), it is plausible that it contributes preferentially to these covert forms of social memory.

To better characterize the involvement of the STN on social memory, we used an optogenetic approach to modulate STN activity during specific phases of social memory processing. These manipulations did not affect object recognition, general social exploration, or social novelty discrimination. In line with lesion data, STN optogenetic inhibition disrupted recognition when applied during either the encoding or recall phase of the social discrimination task. In contrast, optogenetic HF-stimulation impaired memory only when applied during encoding, not recall. Interestingly, only optogenetic inhibition impaired cage-mate recognition. The fact that STN HF photo-inhibition does not impair cage-mate recognition confirms that it only blunts social memory encoding, but not recall. The fact that STN short-term (optogenetic), but not long term (lesions), inhibition blunts rats’ ability to recognize their cage-mate suggest a compensatory mechanism: rats with permanent STN lesions likely adapted over time by relying on alternative sensory cues to identify their cage-mate. In contrast, optogenetic inhibition disrupted STN function acutely during testing, without allowing time for such compensation.

Both optogenetic inhibition and stimulation during the encoding phase impaired recognition, despite being opposite manipulations. This paradox has been reported before (Tiran-Cappello et al., 2018; Vielle et al., 2025) and may be explained by the STN’s integrative role in processing parallel cortical inputs (Parent and Hazrati 1995). Disrupting its activity—regardless of direction—may impair the relay of cortical information to basal ganglia circuits. Alternatively, both manipulations may disrupt local oscillatory dynamics, particularly in the theta band. Theta synchronization has been linked to social novelty detection and memory formation (Buzsáki & Draguhn, 2004, Tendler & Wagner, 2015), and theta power in the STN is modulated by affective salience in PD (Buot et al., 2021; Zénon et al., 2016). Thus, encoding social information may depend on theta activity in the STN, which, once established, could support memory recall unless disrupted by inhibition.

### Conclusions

In sum, our results identify the STN as a key node in the neural network underlying social memory. Its disruption—via lesions or acute optogenetic inhibition—impairs memory for recently encountered peers. HF-stimulation selectively disrupts encoding, but not recall, suggesting a temporally specific role for STN activity. These effects appear independent of general sociability or object memory. Our findings also raise important questions about the functional compensation following chronic STN lesions and the potential consequences of STN modulation in clinical settings. Further investigations, particularly into STN oscillatory dynamics during social memory tasks, are needed to clarify its role in social cognition and its relevance to human neuropathology.

## Data availability statement

Data are available upon request to the corresponding author

## Acknowledgements

The authors thank Alix Tiran-Cappello for his help with optogenetics tool and analyses.

## Authors contribution

CV: designed the experiments, acquired the data, analyzed and interpreted the data and wrote the manuscript **LV:** acquired some data, analyzed and interpreted them, contributed to the manuscript **MW:** acquired some data, analyzed and interpreted them, contributed to the manuscript **NM:** acquired some data, analyzed and interpreted them, contributed to the manuscript **FP and EP:** performed all the histology work **CB:** design ed the experiments, obtained the fundings, interpreted the results wrote the manuscript

## Fundings

CNRS, Aix-Marseille Université, French Ministry of Higher Education, Research and Innovation., IRESP

## Notes

### Competing Interest Statement

The authors have declared no competing interest.

